# Molecular signatures of non-typeable *Haemophilus influenzae* lung adaptation in paediatric chronic lung disease

**DOI:** 10.1101/614214

**Authors:** Ammar Aziz, Derek S. Sarovich, Elizabeth Nosworthy, Jemima Beissbarth, Anne B. Chang, Heidi Smith-Vaughan, Erin P. Price, Tegan M. Harris

**Affiliations:** Child Health Division, Menzies School of Health Research, Charles Darwin University, Tiwi, NT, 0810, Australia; GeneCology Research Centre, University of the Sunshine Coast, Sippy Downs, QLD, 4556, Australia; Department of Respiratory and Sleep Medicine, Children’s Health Queensland, Queensland University of Technology, Brisbane, QLD, Australia

**Keywords:** Non-typeable *Haemophilus influenzae*, NTHi, RNA-seq, transcriptomics, comparative genomics, convergence, bacterial evolution, adaptation

## Abstract

Non-typeable *Haemophilus influenzae* (NTHi), an opportunistic pathogen of the upper airways of healthy children, can infect the lower airways, driving chronic lung disease. However, the molecular basis underpinning NTHi transition from a commensal to a pathogen is not clearly understood. Here, we performed comparative genomic and transcriptomic analyses of 12 paired, isogenic NTHi strains, isolated from the nasopharynx (NP) and bronchoalveolar lavage (BAL) of 11 children with chronic lung disease, to identify convergent molecular signatures associated with lung adaptation. Comparative genomic analyses of the 12 NP-BAL pairs demonstrated that five were genetically identical, with the remaining seven differing by only 1 to 3 mutations. Within-patient transcriptomic analyses identified between 2 and 58 differentially expressed genes in 8 of the 12 NP-BAL pairs, including pairs with no observable genomic changes. Whilst no convergence was observed at the gene level, functional enrichment analysis revealed significant under-representation of differentially expressed genes belonging to *Coenzyme metabolism, Function unknown, Translation, ribosomal structure and biogenesis C*luster of Orthologous Groups categories. In contrast, *Carbohydrate transport and metabolism, Cell motility and secretion, Intracellular trafficking and secretion*, and *Energy production* categories were over-represented. This observed trend amongst genetically-unrelated NTHi strains provides evidence of convergent transcriptional adaptation of NTHi to paediatric airways that deserves further exploration. Understanding the pathoadaptative mechanisms that NTHi employs to infect and persist in the lower paediatric airways is essential for devising targeted diagnostics and treatments aimed at minimising disease severity, and ultimately, preventing NTHi lung infections and subsequent chronic lung disease in children.

## Introduction

Non-typeable *Haemophilus influenzae* (NTHi) is a Gram-negative bacterium that frequently colonises the respiratory tract mucosa of humans. NTHi is part of the upper airway flora and is found in 20-50% of healthy children and 20-30% of healthy adults (Slack, 2015). The upper airways may act as a reservoir for seeding the lung environment (Fothergill et al., 2014), where NTHi switches to an opportunistic pathogenic lifestyle, a transition driven by multiple, poorly understood factors (Duell et al., 2016).

NTHi facilitates the colonisation of other bacterial species in the lower airways (Van Eldere et al., 2014) and is associated with exacerbations in several respiratory diseases (Duell et al., 2016). Indeed, its presence in the lower airways is associated with increased risk of future development of bronchiectasis (Wurzel et al., 2016). In children with chronic suppurative lung disease (CSLD) or bronchiectasis, NTHi is the most commonly detected bacterium in the lower airways (Byun et al., 2017). In adults, the bacterium can chronically and repeatedly colonise the lungs of patients with chronic obstructive pulmonary disease (COPD), bronchiectasis, or COPD with bronchiectasis (Martinez-Garcia et al., 2011; Sriram et al., 2018), resulting in increased airway inflammation (Tufvesson et al., 2015). NTHi also has the capacity to form biofilms (Murphy and Kirkham, 2002; Swords, 2012) resulting in a decreased susceptibility to antibiotics and increased inflammatory and defence responses in the host (Starner et al., 2006). Unsurprisingly, NTHi is a pathogen with increasing public health recognition (Van Eldere et al., 2014; Slack, 2017).

NTHi has evolved several adaptive mechanisms that enable the bacterium to survive and persist in the human host, including modification of its genome via homologous recombination and phase variation (Erwin et al., 2005; Power et al., 2012). Modulation of gene expression through phase variation enables NTHi to rapidly adapt to and colonise new environments (Wong and Akerley, 2012). Analysis of the NTHi genome has identified several phase-variable virulence factors involved in persistence, nutrient uptake, cellular adherence, and host immune response evasion (Pettigrew et al., 2018), with direct modulation of gene expression in response to environmental stimuli and host milieu (Baddal et al., 2015). NTHi is naturally competent and can undergo high rates of recombination, resulting in greater genetic and phenotypic diversity than serotypeable *H. influenzae* (Connor et al., 2012); however, unlike other chronic respiratory infections (Smith et al., 2006; Price et al., 2013), no significant gene loss or gain has yet been observed in chronic *H. influenzae* infections (Pettigrew et al., 2018).

Identifying signals of NTHi adaptation to the lung would aid in unmasking the mechanisms facilitating pathogenesis, which is critical for improving diagnosis, treatment and prevention of NTHi-associated lower airway diseases. The mechanisms involved in the transition from a commensal state in the nasopharynx (NP) to a pathogenic lifestyle in the lower airways is not well understood (De Chiara et al., 2014; Pettigrew et al., 2018). In this study, we investigated genome- and transcriptome-wide differences between 12 isogenic NTHi pairs obtained from the NP and lower airways (via bronchoalveolar lavage; BAL) of patients with bronchiectasis or CSLD. We hypothesised that NTHi isolated from the airways of patients with lung disease would exhibit characteristic genetic or transcriptional profiles that signalled a change from a commensal to a pathogen.

## Methods

### Isolate Collection

NP swabs and BAL specimens were collected concurrently from Australian children (mean, 3.2 years; range: 1.1-13.9 years) diagnosed with bronchiectasis or CSLD (Table 1) at participating hospitals in the Northern Territory and Queensland (Hare et al., 2010). Following collection, specimens were transported on ice in 1 mL skim milk, tryptone, glucose and glycerol broth (STGGB) (Gibson and Khoury, 1986; Hare et al., 2010) and stored at −80°C before processing. All subsequent cultures were also stored in STGGB at −80°C. Following selective culture on chocolate blood agar (CBA) supplemented with bacitracin, vancomycin, and clindamycin (Oxoid, Scoresby, VIC, Australia), NTHi was isolated and confirmed as previously described (Hare et al., 2010; Price et al., 2017). Antibiotic sensitivity was determined by disc diffusion as per the Calibrated Dichotomous Susceptibility (CDS) testing guidelines (8^th^ Edition) (Bell et al., 2016) for ampicillin, amoxicillin-clavulanate, cefaclor, ceftriaxone, chloramphenicol, tetracycline and azithromycin.

**Table 1.** Summary of genomic and transcriptomic differences between paired, isogenic NTHi isolates obtained from the nasopharynx (NP) and lower airways (bronchoalveolar lavage; BAL) of 11 Australian children with chronic lung disease.

| Patient ID | Site of isolation | Isolate no. | No. SNPs | No. indels | No. differentially expressed genes (up/down) | Genes with transcriptional and genetic alterations |
| --- | --- | --- | --- | --- | --- | --- |
| 60051 | NP | 1 | 0 | 1 | 58 (4/54) | *hsdM3* |
|  | BAL | 1 |  |  |  |  |
| 60068 | NP | 3 | 0 | 1 | 54 (1/53) | --- |
|  | BAL | 1 |  |  |  |  |
| 60069 | NP | 2 | 0 | 1 | 0 | --- |
|  | BAL | 1 |  |  |  |  |
| 60294 | NP | 1 | 0 | 0 | 0 | --- |
|  | BAL | 1 |  |  |  |  |
| 60295 | NP | 1 | 1 | 0 | 46 (38/8) | --- |
|  | BAL | 1 |  |  |  |  |
| 60316 | NP | 1 | 0 | 0 | 0 | --- |
|  | BAL | 1 |  |  |  |  |
| 60362 | NP | 1 | 0 | 0 | 15 (14/1) | --- |
|  | BAL | 2 |  |  |  |  |
| 60370 | NP | 2 | 0 | 0 | 2 (2/0) | --- |
|  | BAL | 2 |  |  |  |  |
| 60373_P1 | NP | 1 | 0 | 1* | 5 (3/2) | *hsdM3* |
|  | BAL | 1 |  |  |  |  |
| 60373_P2 | NP | 3 | 0 | 0 | 4 (4/0) | --- |
|  | BAL | 4 |  |  |  |  |
| 65001 | NP | 2 | 0 | 3 | 0 | --- |
|  | BAL | 1 |  |  |  |  |
| 65290 | NP | 3 | 1 | 0 | 4 (4/0) | --- |
|  | BAL | 4 |  |  |  |  |

Abbreviations: SNP, single-nucleotide polymorphism; indel, insertion-deletion.
\*The BAL strain from this patient contains a mixture of two deleterious indel mutations in *hsdM3* at a 40:60 ratio (See Figure 1).

### Whole-Genome Sequencing and Comparative Genomics

DNA extraction was performed as previously described (Price et al., 2015). Whole-genome sequencing (WGS) was carried out at the Australian Genome Research Facility (Parkville, VIC, Australia) using the Illumina (San Diego, CA, USA) HiSeq 2500 platform. As a criterion for our study, all isolate pairs were required to be highly genetically similar (i.e. isogenic). To identify isogenic NP-BAL pairs from each patient, multilocus sequence typing (MLST) (Jolley and Maiden, 2010) followed by phylogenetic analysis was performed. Further details on isolate selection is described in Text S1. Based on this analysis, we selected 12 paired NP-BAL isolates from 11 patients for further investigation; two genetically distinct NP-BAL pairs were selected from one patient (designated 60373_P1 and 60373_P2).

To catalogue the mutations separating each NP-BAL pair, SPANDx v3.2 (default settings) (Sarovich and Price, 2014) was used. Where possible, genetic variants were also confirmed using transcriptomic (RNA-seq) data. For each analysis, the NP isolate draft assembly was used as the reference for variant calling. Further details are provided in Text S1.

### NTHi Liquid Media Growth Conditions for RNA Harvest

Due to the reliance of NTHi growth on the presence of X (haemin/haematin) and V (nicotinamide-adenine-dinucleotide) blood factors (Holt, 1962), we initially attempted to culture isolates in brain heart infusion (BHI) broth supplemented with Haemophilus Test Medium (HTM; Oxoid), and subsequently, a HTM-supplemented artificial sputum medium formulated to mimic the mucosal environment in the airways of cystic fibrosis patients (Fung et al., 2010; Price et al., 2017). However, due to inconsistent optical density (OD) readings and poor growth or flocculation of some or all isolates in these media, we investigated an alternative liquid medium for the growth of NTHi for RNA-seq (see Text S1 for further details). CBA is widely used for culturing NTHi and other fastidious bacteria (Artman et al., 1983). Thus, we used this medium as the basis for formulating liquid chocolate medium (LCM). Formulation of LCM and growth conditions are provided in Text S1.

### RNA Extraction

All isolates were cultured in LCM and in duplicate to account for any technical replicability issues. RNA was extracted using TRIzol® as described in Text S1. After DNA digest, absence was verified using real-time PCR assays that detect Hi DNA (Aziz et al., 2017) and defibrinated horse blood DNA (Humair et al., 2007). For further details see Text S1.

### Differential Expression (DE) Analysis

RNA-seq processing and DE analysis was performed as described in Text S1. Breifly, RNA-seq was carried out using the Illumina HiSeq 2500 platform. In total, 24 isolates were sequenced in duplicate, generating 48 transcriptomes.

To identify molecular signatures of lung adaptation on a within-patient basis, DE analysis was first conducted by comparing BAL with NP isolates from individual patients. Subsequently, convergence analysis was performed by comparing BAL with NP isolates from all 12 isolate pairs. The following additional analyses with different sample sets were performed to ensure robustness: a) those NP-BAL pairs with DE detected in the within-patient analysis (eight in total), and b) the three NP-BAL pairs with >45 DE genes (60051, 60068,60295). For all convergence analyses, the two NP-BAL pairs from patient 60373 were treated either as independent NP-BAL pairs or grouped as a single patient. DE was conducted using edgeR (v3.18.1) (Robinson et al., 2010), implemented in R (v3.4.1) (R Core Team, 2014). For all analyses, DE was determined using the quasi-likelihood method (*glmQLFit* function).

Genes with significant DE were annotated using Clusters of Orthologous Groups (COG) (Galperin et al., 2015), with categories retrieved from a previous study (Santana et al., 2014) with minor corrections (Table S1). For genes with multiple COG categories, all assigned categories were included in the analysis. To assess for significant enrichment of functional groups, COG categories of the reference genome 86-028NP were compared with those identified in the within-patient DE analyses. Statistical assessment comparing the proportion of categories was performed in R using a two-tailed Fisher’s exact test with a false discovery rate corrected *p*-value of ≤0.05.

### Data Availability

All genomic and transcriptomic data generated as part of this study are publicly available under NCBI BioProject PRJNA484075. Genome assemblies for the 24 NTHi strains can be found in NCBI GenBank (QWLW00000000-QWMT00000000), and Illumina WGS and RNA-Seq data are available on the Sequence Read Database (SRR7719256-SRR7719303).

## Results

### Comparative Genomic Analysis of Paired Isolates

To assess diversity and to identify potential clustering of NP or BAL isolates, a maximum parsimony phylogenetic tree was constructed using 124,262 biallelic, core-genome SNPs identified among the paired isolates and a global NTHi isolate set comprising 157 strains (Figure S1). Phylogenomic analysis confirmed that pairs from individual patients clustered closely together yet were distinct from other NP-BAL pairs. Consistent with previous studies (De Chiara et al., 2014; Price et al., 2015; Pettigrew et al., 2018), there was no evidence of NP-or BAL-specific clades. Isolate pairs were identical sequence types (STs), with 5 of the 12 pairs possessing novel STs (ST-1906 to ST-1910; Table S2). Patient 60373 had two distinct isolate pairs (NP-BAL pairs 60373_P1 and 60373_P2) that were separated by >17,000 SNPs, demonstrating the presence of at least two NTHi lineages colonising this patient. This finding was also reflected in the MLST data, with 60373_P1 isolates belonging to ST-1910 whereas 60373_P2 isolates were ST-139 (Table S2). Antibiotics are routinely used to treat exacerbation events in paediatric chronic lung disease, and resistance to clinically-relevant antibiotics can arise during such infections. In the current work, no NTHi isolate exhibited resistance towards any of the tested antibiotics, ruling out antibiotic resistance-driven mutations arising between isolate pairs.

Next, within-patient comparative genomic analyses were performed to identify variants separating the paired NP-BAL isolates. Pairs were highly genetically similar, with a maximum of three mutations observed between any one pair (Table 1). In total, two non-synonymous SNPs and seven indels (six in open-reading frames) were identified among seven NP-BAL pairs, with the remaining five NP-BAL pairs being genetically identical (Table 2). No copy-number variants, large deletions or inversions were observed between any of the pairs. One non-synonymous SNP occurred in the exogenous haem acquisition-encoding gene, *hgpB* (HgpB_Arg407Lys_; 60295 BAL Hi1), and the second affected the outer membrane protein assembly factor, *bamA* (BamA_Glu508Gly_; 65290 NP Hi3). Of the seven indels, four were simple sequence repeats (SSRs) that were predicted to result in frameshift mutations, leading to truncated proteins in phase-variable genes (HsdM3_Glu9fs_, HsdM3_Glu12fs_, HgpB_Thr35fs_, and Lic3A_Ser18fs_ in 60051 BAL Hi1, 60373_P1 BAL Hi1, 60068 BAL Hi1, and 65001 BAL Hi1, respectively). The remaining three indels were in: *ompP2* (OmpP2_Ala269_Gly270insValGlyAla_; 65001 BAL Hi1), *hsdS2* (HsdS2_Thr173_Leu176del_; 65001 BAL Hi1), and an intergenic region 62bp upstream of *hxuC* (60069 BAL Hi1). Notably, *hsdM3* (in 60051 and 60373_P1) and *hgpB* (in 60068 and 60295) were mutated in isolates from two patients. When compared with 86-028NP, all mutations were shown to deleteriously affect the BAL isolates (Table 2).

**Table 2.**
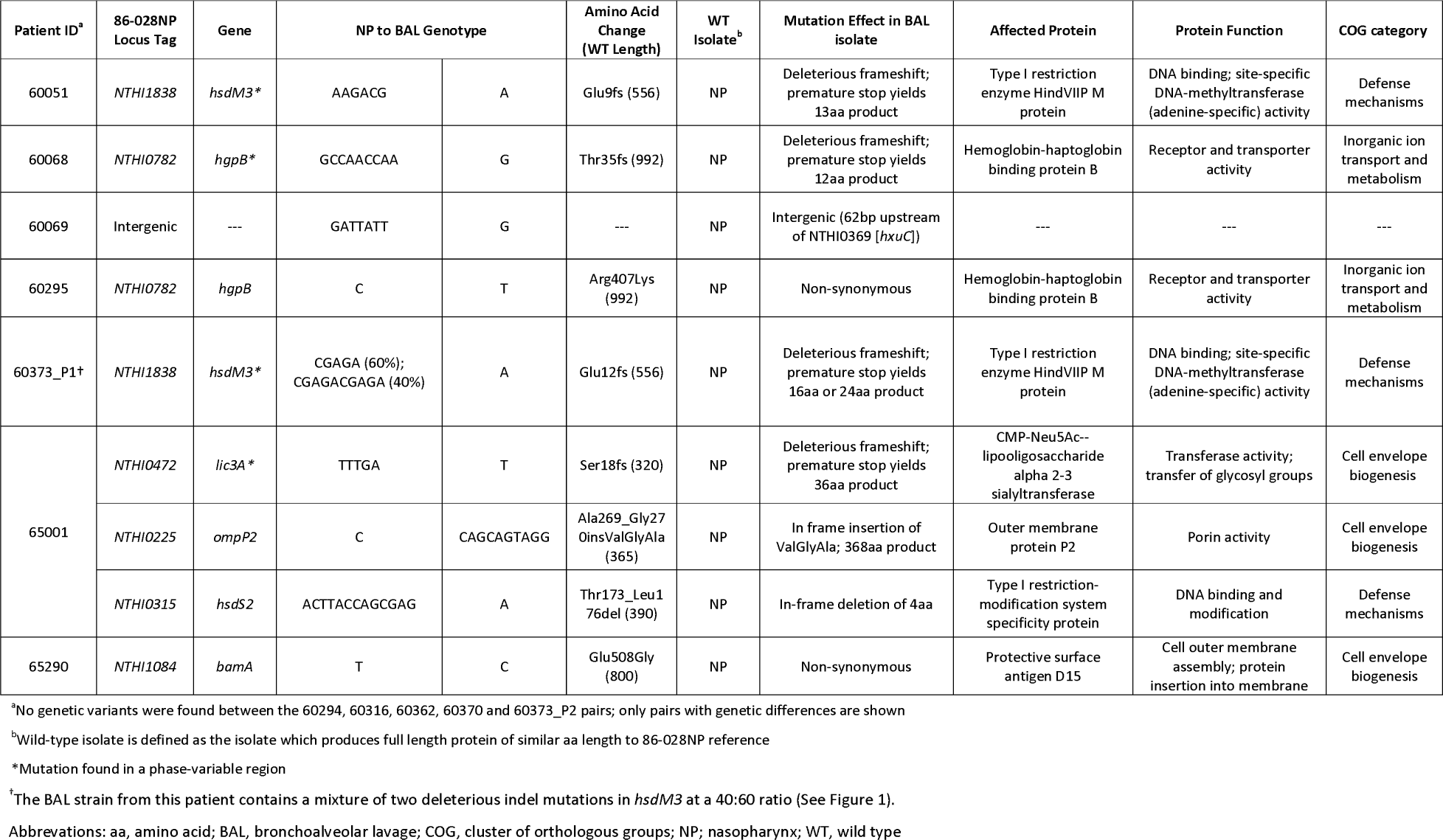
Summary of genetic variants identified between 12 paired, isogenic non-typeable *Haemophilus influenzae* isolates retrieved from the sopharynx vs. lower airways (bronchoalveolar lavage; BAL).

The *hsdM3* indels in both the 60051 and 60373_P1 BAL isolates occurred in the same pentanucleotide SSR tract, which comprises variable copy number of a ‘CGAGA’ motif at the 5’ end of this gene. Phase variation in *hsdM3* is driven by changes in this highly mutable SSR tract and is influenced by a lack of adenine methylation (Zaleski et al., 2005). A 5bp AGACG deletion in this SSR tract in 60051 BAL Hi1 (Figure 1A and 1C) is predicted to produce a truncated protein of just 13 residues. Similarly, a 5bp CGAGA deletion was observed in 60373_P1 BAL Hi1; however, it was only supported by ∼60% of aligned reads. Upon closer inspection, a 10bp deletion (CGAGACGAGA) was present in the remaining ∼40% of aligned reads (Figure 1B). This mixture of *hsdM3* alleles is supported by the RNA-seq reads. The 5bp (∼60%) and 10bp (∼40%) components were both predicted to produce a truncated protein of 16 and 24 residues, respectively (Figure 1C). Given that isolates were subjected to a genetic bottleneck prior to sequencing, these mixtures suggest that the *hsdM3* SSR undergoes a rapid mutation rate, with mutations potentially occurring during laboratory passage.

**Figure 1.**
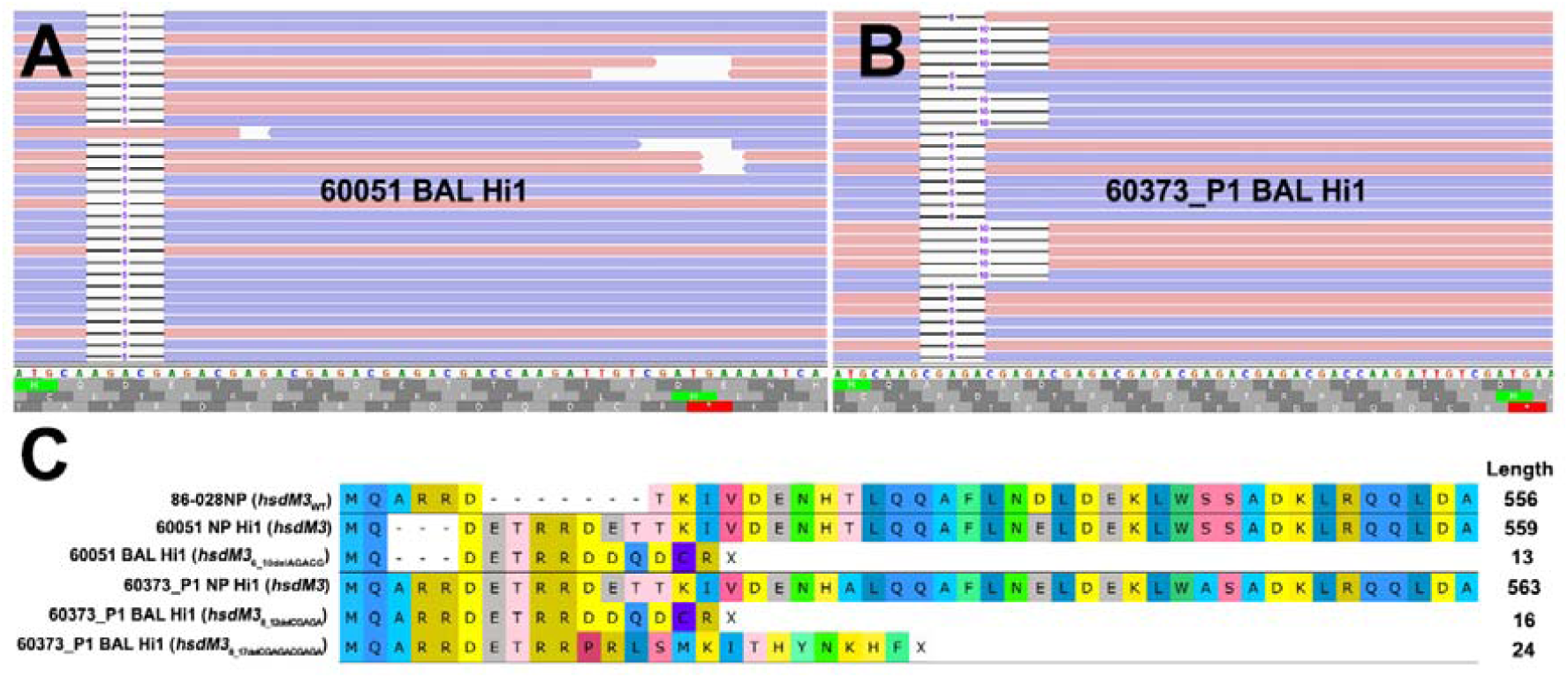
Read mapping analysis of the highly mutable *hsdM3* pentanucleotide short sequence repeat (SSR) tract in 60051 and 60373_P1 NP-BAL pairs. (A) Graphical representation of 60051 BAL Hi1 illumina reads aligned against the *hdsM3* region of the 60051 NP Hi1 reference assembly. One allele consisting of a 5bp deletion (AGACG) was observed. (B) Graphical representation of 60373_P1 BAL Hi1 illumina reads aligned against the 60373_P1 NP Hi1 reference assembly. Two different alleles consisting of 5bp (AGACG) and 10bp (AGACGx2) deletions were observed in the BAL isolate. (C) Amino acid alignment of *hsdM3* against reference strain 86-028NP (wild-type). NP isolates from 60051 and 60373_P1 encode the full-length protein, with minor differences resulting from the variable SSR track. Indels in the same SSR track of both BAL isolates, including the two alleles observed in 60373_P1, result in a deleterious frameshift that is predicted to yield a truncated protein of 13-24aa.

### Liquid Chocolate Medium for Reproducible NTHi Growth

As NTHi has an absolute growth requirement for X and V blood factors (Holt, 1962), commonly used culture media for this fastidious bacterium typically includes lysed blood in a nutrient-rich agar base (e.g. CBA) (Artman et al., 1983), or BHI base supplemented with HTM (Coleman et al., 2003). However, we found uniform and reproducible growth of NTHi in HTM-supplemented BHI problematic; variable growth rates were observed both among isolates and between technical replicates, indicating inconsistent bacterial growth. Additionally, the microbial load was insufficient for RNA-seq, even after 24h incubation. We also observed inconsistent OD readings between replicates and a tendency towards flocculation in certain strains. Similar issues were observed with HTM-supplemented artificial sputum medium, which is designed to mimic cystic fibrosis sputum (Fung et al., 2010) (Figure S2A).

To overcome these issues, we developed liquid chocolate medium (LCM), a medium that is straightforward to formulate, inexpensive and produces high-quantity NTHi cellular density for downstream analysis. LCM yielded NTHi cultures with reproducible growth, consistent growth rates, little to no flocculation, and sufficient microbial load to support sequencing of late-log phase RNA. LCM was thus used to determine the growth curves of four isolates to identify an appropriate culture duration for RNA extraction. A 7.5-hour post-inoculation time point was found to represent late-log phase growth in all four cultures tested (Figure S2B). This time point, which was chosen to reflect non-logarithmic bacterial growth rates *in vivo* (Yang et al., 2008; Kasi et al., 2017), was subsequently used for RNA extraction of all isolates.

### Within-patient DE and Comparison with Genetic Mutations

Culture and RNA extraction of 24 isolates was performed in duplicate. On average, 89.2% (87.1-92.0%) of sequence reads aligned uniquely to the 86-028NP reference genome, and of these, an average of 70% (68.2-74.8%) were assigned to known genomic features (Table S2). Due to this high mapping percentage, within-patient DE analysis was performed using 86-028NP as the reference genome. This approach also greatly simplified comparisons among patients. DE was observed between eight NP-BAL pairs, ranging from 2 to 58 DE genes per pair. Three pairs (60051, 60068, and 60295) had >45 DE genes, five pairs (60362, 60370, 60373_P1, 60373_P2, and 65290) had 15 or fewer DE genes (Dataset S1), and the remaining four pairs (60069, 60294, 60316, and 65001) did not exhibit any DE.

In total, 189 DE genes, representing a wide functional spectrum, were detected across the eight NP-BAL pairs. Eight genes were DE in three pairs (*comE, comM, glgC, ligA, pilB, pilC*, smf and *tnaA*), 39 were DE in two pairs (*arcC, argF, asnA, comABCDF, dmsC, glgABPX, glpK, hitA, hsdMS3, licBC, mao2, nagAB, nanAEK, NTHI0229, NTHI0232, NTHI0235, NTHI0646, NTHI0647, NTHI1183, NTHI1243, nrfCD, pckA, pilA, sdaA, siaT*, and *tnaB*), and the remaining 87 were DE in single NP-BAL pairs (Figure 2; Dataset S1). However, only ten genes exhibited concordant downregulation (*mao2, NTHI0646, NTHI0647, pckA, sdaA*) or upregulation (*asnA, hitA, hsdMS3, hsdS3, tnaB*) in the BAL isolates. In all other DE genes, discordance in the directionality of expression was observed, including the eight shared DE genes in three NP-BAL pairs (Figure 2). Additionally, genes in the 60295 pair were predominantly upregulated, whereas genes were predominantly downregulated in pairs 60051 and 60068. Overall, there was little overlap amongst patients.

**Figure 2.**
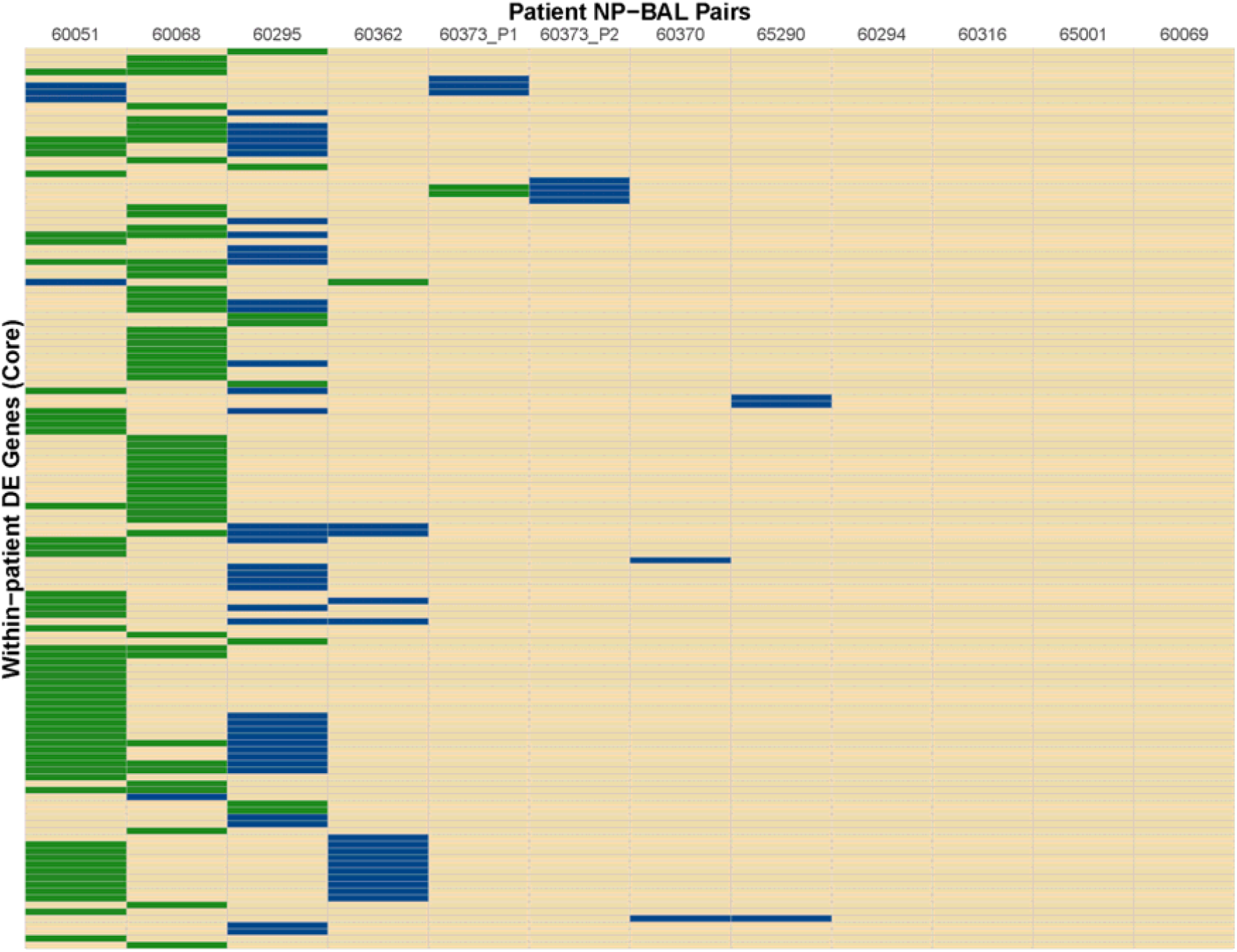
Heat map of differentially expressed (DE) genes identified between the NP-BAL pairs from 11 paediatric chronic lung disease patients. In total 134 non-redundant DE genes were detected across eight NP-BAL pairs. DE was observed in eight NP-BAL pairs; the remaining four pairs contained no DE. Genes were predominantly upregulated in the 60295 NP-BAL pair but predominately downregulated in the 60051 and 60068 NP-BAL pairs. The direction of log_2_ fold change of DE genes in BAL is colour-coded as follows: upregulation (blue), downregulation (green), no DE (tan).

There was also minimal correlation between the number of genetic mutations and the number of DE genes detected between NP-BAL pairs (Table 1). For example, the 60295 NP-BAL pair exhibited 46 DE genes, but only a single non-synonymous SNP (HgpB_Arg407Lys_) was observed in the BAL isolate. In contrast, no DE was detected between the 65001 NP-BAL pair (Dataset S1), despite three indels being identified in the BAL isolate, including a deleterious frameshift in the phase-variable sialyltransferase gene, *lic3A*, which truncates the Lic3A protein from 320 to just 36 residues (Table 2). Two NP-BAL pairs (60294 and 60316) contained no mutations and no DE genes. Conversely, two NP-BAL pairs (60051 and 60373_P1) contained both genetic and transcriptional alterations affecting a single phase-variable gene, *hsdM3* (Table 2; Dataset S1). For both pairs, *hsdM3* and *hsdS3* were upregulated in the BAL isolates, with an average log2 fold change of 2.5 and 4.6, respectively. Additionally, *hsdR3* was upregulated by 2.2-fold in 60373_P1. However, both BAL isolates from 60051 and 60373_P1 encode truncated HsdM proteins of 13 and 16-24 residues, respectively (Figure 1C), indicating that *hsdM3* upregulation in these isolates is unlikely to have a functional consequence. Notably, the two isolates with mutations in *hgpB*, 60068 BAL Hi1 and 60295 BAL Hi1, both had a relatively high amount of DE, with 55 and 46 DE genes, respectively.

### DE Convergence and Functional Enrichment Analysis of NP-BAL Isolates

We compared BAL with NP isolates from all patients to identify genes with convergent gene expression. However, no single convergent genetic mutations or DE genes were identified, suggesting more complex pathways involved in lung adaptation. Therefore, we conducted a functional enrichment analysis of the 134 non-redundant DE genes using COG assignments to identify enriched categories. Statistical analysis identified 4/21 COG categories that were significantly over-represented in the functional enrichment analysis (*Carbohydrate transport and metabolism, Energy production and conversion, Intracellular trafficking and secretion*, and *Cell motility and secretion*), and 3/21 categories (*Translation, ribosomal structure and biogenesis, Coenzyme metabolism*, and *Function unknown*) that were under-represented (*p*≤0.05, Figure 3). Of note, the *Function unknown* category encompasses 20.2% genes in the 86-028NP genome, yet DE was only observed in 3.5% of these genes. The basis for this under-representation may partly reflect the artificially high number of gene annotations in the NTHi genome. In support of this notion, 47/397 (11.8%) genes assigned to the *Function unknown* category exhibited zero reads across our 48 transcriptomes (Dataset S2, grey shaded rows), representing 37.9% of all genes with zero reads. Approximately 51.8% of the other genes with zero reads corresponded with paralogous ribosomal and transfer RNA species (Dataset S2).

**Figure 3.**
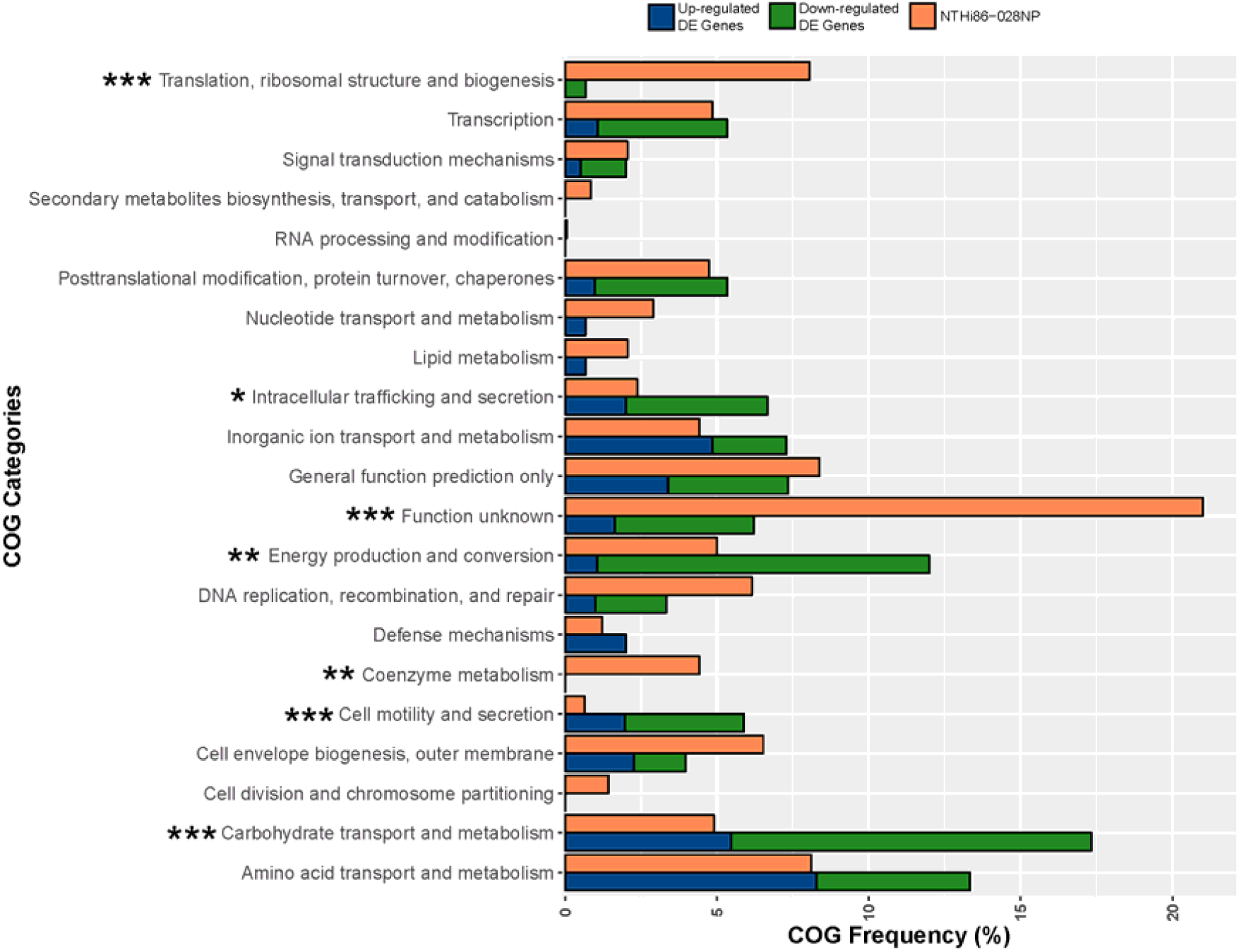
Clusters of Orthologous Group (COG) analysis comparing the frequency of categories in the 86-028NP genome with all DE genes in the 12 NP-BAL pairs. Seven COG categories were significantly enriched in DE genes from BAL-derived isolates, comprising four over-represented categories *(Carbohydrate transport and metabolism, Cell motility and secretion, Energy production and conversion* and *Intracellular trafficking and secretion*) that were majority downregulated and three under-represented categories *(Coenzyme metabolism, Function unknown*, and *Translation, ribosomal structure and biogenesis). *** p* <0.001, ** p< 0.01, *p< 0.05.

The most significant over-represented COGs (*p* < 0.001) were *Carbohydrate transport and metabolism*, which is represented by 4.9% of 86-028NP genes, yet 17.3% of DE genes belonged to this category, and *Cell motility and secretion*, with only 0.6% genes in 86-028NP but with 5.8% of DE genes belonging to this COG. The most significant under-represented COGs were *Translation, ribosomal structure and biogenesis*, comprising 8.0% of 86-028NP genes but only 0.6% of the DE genes, and *Function unknown*, which comprises 20.2% of 86-028NP genes yet only 6.2% of DE genes belonged to this category (Figure 3; Dataset S1). Other significant COG categories (*p* < 0.01) were *Energy production and conversion* (5.0% of 86-028NP genes but 12.0% DE genes) and *Coenzyme metabolism* (4.4% 86-028NP genes but 0% DE genes).

## Discussion

NTHi is a pathogen of emerging public health importance (Erwin and Smith, 2007; Van Eldere et al., 2014). Conventionally considered a commensal of the upper airways, it is now recognised that NTHi is associated with increased severity and progression of multiple polymicrobial diseases of the lower airways such as COPD, bronchiectasis, and CSLD (Murphy, 2003; Purcell et al., 2014). In this study, we aimed to identify signals of NTHi pathoadaptation to the lower airways using 12 paired isogenic strains retrieved from the upper (NP) and the lower (BAL fluid) airways of 11 paediatric patients with CLSD or bronchiectasis. Similar studies have employed WGS to document the genome-wide changes leading to chronic adaptation and persistence of NTHi retrieved from COPD sputum in adults (Moleres et al., 2018; Pettigrew et al., 2018). The current work expands on these prior studies by examining both the genomic and transcriptomic profiles of NTHi adaptation in children’s airways. To ensure isolates were retrieved from the lower airways, we obtained NTHi isolates from the lower airways (BAL fluid) rather than sputum, the latter of which is fraught with upper respiratory tract contamination issues (Grønseth et al., 2017).

As part of our study criteria, only isogenic strains were selected to simplify the identification of convergent pathoadaptative mechanisms. As expected, comparative genomic analyses found minimal genetic differences between all isogenic NP-BAL pairs, with only nine total variants in our dataset, ranging from zero to three variants per pair. Five pairs were genetically identical, with the remaining seven encoding a total of two SNPs and seven indels. Notably, four of these indels occurred in tandem repeat regions, or SSR tracts, of phase-variable genes, which are abundant in the NTHi genome (Gilsdorf et al., 2004; Pettigrew et al., 2018). SSRs provide a rapid mechanism for gene expression modulation and adaptation (Vasu and Nagaraja, 2013; De Ste Croix et al., 2017). Two SSR indels occurred in the hypervariable Type I restriction-modification (RM) system methyltransferase gene, *hsdM3.* The remaining SSR indels occurred in the haemoglobin-haptoglobin binding protein B-encoding gene *hgpB*, which is involved in NP colonisation and iron acquisition (Poole et al.), and *lic3A*, which encodes a lipooligosaccharide (LOS) sialyltransferase that facilitates NTHi colonisation in different anatomical sites (Hosking et al., 1999; Hood et al., 2001).

Our genetic findings concur with the recent studies, which found that phase variation in *lic3A, hsdM3* and *hgpB* are common in NTHi isolates retrieved from COPD airways (Moleres et al., 2018; Pettigrew et al., 2018). In addition to the *hgpB* SSR variant in 60068 BAL Hi1 that truncates the protein from 992 to 12 residues, a non-synonymous SNP (HgpB_Arg407Lys_) was seen in 60295 BAL Hi1. Likewise, the SSR region of *hsdM3* was deleteriously mutated in the 60051 and 60373_P1 BAL isolates, yielding truncated proteins of ≤24 residues (Figure 1C). The *hsdM3* gene is part of a Type I RM system that acts as a defence system against foreign DNA invasion by, e.g. bacteriophages (Murray, 2000), fine-tuning of competence through variable methylation patterns, and in the methylation of self-DNA that may alter gene expression (Vasu and Nagaraja, 2013). While the *hdsM3* mutations in the paediatric BAL isolates would render their corresponding Type I RM systems non-functional, the phase-variable nature of *hsdM3* enables rapid reverse switching back to an active state. Others have shown that phase-variable RM systems allow NTHi to rapidly generate genomic and phenotypic diversity, which may enhance survival and persistence, especially in new environments (Zaleski et al., 2005; Atack et al., 2015). Taken together, the commonality of SSR tract mutants affecting *hgpB, hsdM3* and l*ic3A* in airway-derived isolates demonstrates that phase variation of these loci is driven by strong diversifying selection, with this variation being key in the long-term persistence of NTHi within the hostile lung environment.

Outer membrane porin (OmpP1 and OmpP2) mutations are also common in COPD-derived NTHi isolates. Previous studies have identified deleterious *ompP1* mutations in ∼33% of COPD isolates, and in-frame *ompP2* mutations (i.e. missense SNPs and indels) in ∼20% of COPD isolates (Moleres et al., 2018; Pettigrew et al., 2018). Similarly, we identified an in-frame *ompP2* mutation in the 65001 BAL Hi1 isolate that occurred in the extracellular loop six motif (Sikkema and Murphy, 1992) and increased OmpP2 length by three amino acids. However, we did not observe any genomic or transcriptomic *ompP1* variation between our 12 NP and BAL pairs. Although our sample size is modest, this result suggests that *ompP1* variants may be much less common in paediatric NTHi lung-adapted strains than COPD lung-adapted strains. Biochemical profiles in COPD airways (e.g. fatty acids with bactericidal activity against NTHi wild-type strains) (Moleres et al., 2018) are potentially different to those present in paediatric bronchiectasis or CSLD airways, which may explain the greater selective pressure for OmpP1 inactivation in COPD airways. Further work is needed to document the prevalence of *ompP1* mutations in a larger paediatric lung isolate cohort to examine whether diseased paediatric airways do in fact impart different OmpP1 pressures on NTHi compared with isolates from COPD airways.

Across the 12 within-patient DE analyses, a total of 134 non-redundant genes (range: 0 to 58) were DE between NP-BAL pairs, with no correlation between the number of mutations and the number of DE genes. Indeed, NP-BAL pairs 60362, 60370 and 60373_P2 were devoid of mutations, yet DE was observed in 15, 2, and 4 genes, respectively. Likewise, 65001 BAL Hi1 encoded mutations in *lic3A, ompP2* and a Type I RM system specificity gene, *hsdS2*, yet no DE was detected in this strain. This lack of correlation suggests that DE between isolate pairs is influenced by epigenetic factors (e.g. altered DNA methylation (Fox et al., 2007)), or by unidentified genetic mutations due to inherent limitations with short-read sequencing approaches (e.g. insertion sequence-mediated rearrangements or variants in paralogues). Future studies employing methylation signature detection and long-read sequencing will enable a deeper understanding of the basis for DE in these strains.

We did not identify any DE gene(s) common to all isolates, indicating no universal expression profile associated with lung adaptation (i.e. the BAL isolates). However, some degree of gene convergence was observed, with eight DE genes common to three isolate pairs, and 39 common to two pairs (Dataset S1). Of these, the *com* and *pil* operons, required for the biogenesis and function of the type IV pilus, were DE in three pairs. The type IV pilus is involved in multiple biological roles including adherence to epithelial cells, competence, twitching motility, biofilm formation and long-term NP colonisation (Bakaletz et al., 2005; Jurcisek et al., 2007). The pathway responsible for uptake, transport and incorporation of N-Acetylneuraminic acid (Neu5Ac) into LOS (Apicella, 2012; Wong and Akerley, 2012), was DE in two pairs. Incorporation of Neu5Ac into the LOS enhances immune evasion and increases resistance to killing by human serum (Mandrell et al., 1992). Most DE genes (*n*=87) occurred in single pairs, suggesting multiple, parallel strategies towards lower airway adaptation. In addition, the directionality of DE was inconsistent, with gene upregulation in some pairs and downregulation in others (Figure 2). Lack of a convergent transcriptional signal across all strains may be due to sampling difficulties, for example, the inclusion of BAL isolates that have yet to adapt to the lower airways or re-seeding of the NP environment with lung-adapted strains. Greater breadth and depth of strains is needed to determine whether a convergent signal of lung adaptation can be identified in the NTHi transcriptome.

To identify parallel signatures of adaptation to the lung environment on a broader scale, we categorised the functions of within-patient DE genes based on COG categories. Overall, strains isolated from the lower airways showed significant enrichment of DE genes in 7/21 COG categories. Four categories were over-represented in the DE gene dataset when compared with their prevalence in the 86-028NP genome, with *Carbohydrate transport and metabolism, Cell motility and secretion, Intracellular trafficking and secretion*, and *Energy production and conversion* being significant (*p* <0.05 or *p* <0.001). In comparison, the COG categories *Translation, ribosomal structure and biogenesis, Function unknown* and *Coenzyme metabolism* were significantly under-represented (*p* <0.01 or *p* <0.001) in the DE gene dataset, suggesting that altered expression of these gene categories is negatively associated with lung adaptation. However, analysis of the 48 transcriptomes revealed that 11.8% genes in the *Function unknown* category had zero reads mapping to the 86-028NP genome (Dataset S2, grey-shaded rows), suggesting that many of these so-called genes likely do not produce messenger RNA or proteins under any growth conditions, and may represent incorrect gene annotations. Further RNA-seq studies of NTHi cultures grown under different conditions will help to resolve these potential annotation issues. Another noteworthy observation was that *Carbohydrate transport and metabolism* and *Energy production and conversion* collectively contained ∼35% of all DE genes and were primarily downregulated. This observation is consistent with Baddal and colleagues, who observed the downregulation of genes involved in NTHi central metabolism in response to environmental stimuli (Baddal et al., 2015). *Intracellular trafficking and secretion* and *Cell motility and secretion* were also over-represented in the BAL isolates and predominately downregulated (Figure 3; Dataset S1). Taken together, these results suggest that NTHi alters certain functional pathways in response to the lower airway environment, suggesting a degree of predictability in adapting to this new niche. Although further work is needed to consolidate these findings, our work sets the stage for identifying key functional pathways that may be exploited for targeted treatment and eradication of this pathogen.

Our study chose not to assess the wider polymicrobial community of the lower airways, as accurate transcriptional characterisation of entire bacterial communities remains bioinformatically challenging (Lim et al., 2013). We instead chose to focus on an in-depth genomic and transcriptional analysis of NTHi, a species known to play a crucial role in the progression and severity of lung disease (Erwin and Smith, 2007; Van Eldere et al., 2014; Slack, 2017). Genetically diverse NTHi isolates from different patients were selected to enable identification of convergent adaptation strategies to the lung environment of paediatric patients. Due to the necessary passaging of strains to ensure purity and to obtain sufficient RNA material for sequencing, we cannot discount that these procedures may have influenced the NTHi expression profiles so that they no longer represent their *in vivo* counterparts. However, as previously demonstrated with other bacterial species, isolates appear to maintain their expression profiles even after microbiological handling and culturing *in vitro* (Viberg et al., 2017). For future studies, we recommend maximising the number of strains isolated from individual patients together with longitudinal sampling to better understand pathogen adaptation over time.

In conclusion, our study provides new insights into NTHi adaptation to the lower airways in paediatric chronic lung diseases. Through genomic and transcriptomic characterisation of 12 paired NTHi isolates from the NP and the lung, we found that NTHi employs several avenues of pathoadaptation that enables this pathogen to persist in the lower airways. Although we did not identify evidence of convergent pathoadaptive mechanisms at the single-gene level, our study identified parallel DE at the functional level, suggesting that NTHi adaptation to the paediatric airways is a complex but ultimately predictable process. The next steps require characterising larger isolate panels using genomic and transcriptomic methods to better understand the pathoadaptive mechanisms that enable NTHi to persist and cause disease. Such findings will be crucial for the informed development of effective therapeutic interventions to prevent NTHi-driven chronic lung diseases

## Author Contributions

AC conducted patient recruitment and specimen collection. EN, JB conducted specimen processing, genotyping, and DNA extractions. AA, JB, HSV, EPP, TMH conducted isolate selection. AA conducted culturing, RNA extraction, and bioinformatics analysis with supervision and assistance from DSS, EPP, TMH. AA wrote the initial manuscript draft, and DSS, EPP and TMH critically reviewed and edited the manuscript. DSS, HSV, AC, and EPP conceived of the study and DSS, HSV, and EPP obtained funding. All authors reviewed and approved the final manuscript.

## Acknowledgements

We wish to thank the Menzies School of Health Research Child Health team for assistance with microbiological aspects of this study, and Roisin Ure (NHS Greater Glasgow and Clyde, Glasgow, UK) for assistance with *H. influenzae* MLST database submission.

## Supplemental material legends

**Text S1.** Supplementary methods.

**Figure S1.** Maximum parsimony phylogenomic analysis of non-typeable *Haemophilus influenzae* isolates from paediatric lung disease against a global strain dataset.

**Figure S2.** Non-typeable *Haemophilus influenzae* growth kinetics in different growth media.

**Table S1.** Updated Cluster of Orthologous Group terms for non-typeable *Haemophilus influenzae* strain 86-028NP. Figshare link: https://doi.org/10.6084/m9.figshare.7956191.v1

**Table S2.** Whole-genome sequencing and RNA sequencing summary information. Figshare link: https://doi.org/10.6084/m9.figshare.7956194.v5

**Dataset S1.** Within-patient Non-typeable *Haemophilus influenzae* differential expression analysis comparing nasopharynx against bronchoalveolar isolates. Figshare link: https://doi.org/10.6084/m9.figshare.7956158.v1

**Dataset S2.** RNA sequencing raw read counts against Non-typeable *Haemophilus influenzae* strain 86-028NP. Figshare link: https://doi.org/10.6084/m9.figshare.7956161.v1

## References

Apicella, M.A. (2012). Nontypeable Haemophilus influenzae: the role of N-acetyl-5-neuraminic acid in biology. Nontypeable Haemophilus influenzae: the role of N-acetyl-5-neuraminic acid in biology 2, 19. doi:10.3389/fcimb.2012.00019.

Artman, M., Domenech, E., and Weiner, M. (1983). Growth of Haemophilus influenzae in simulated blood cultures supplemented with hemin and NAD. Growth of Haemophilus influenzae in simulated blood cultures supplemented with hemin and NAD 18(2), 376–379.

Atack, J.M., Srikhanta, Y.N., Fox, K.L., Jurcisek, J.A., Brockman, K.L., Clark, T.A., et al. (2015). A biphasic epigenetic switch controls immunoevasion, virulence and niche adaptation in non-typeable Haemophilus influenzae. Nat Commun 6. doi:10.1038/ncomms8828.

Aziz, A., Sarovich, D.S., Harris, T.M., Kaestli, M., McRobb, E., Mayo, M., et al. (2017). Suspected cases of intracontinental Burkholderia pseudomallei sequence type homoplasy resolved using whole-genome sequencing. Suspected cases of intracontinental Burkholderia pseudomallei sequence type homoplasy resolved using whole-genome sequencing 3(11). doi:10.1099/mgen.0.000139.

Baddal, B., Muzzi, A., Censini, S., Calogero, R.A., Torricelli, G., Guidotti, S., et al. (2015). Dual RNA-seq of Nontypeable Haemophilus influenzae and Host Cell Transcriptomes Reveals Novel Insights into Host-Pathogen Cross Talk. Dual RNA-seq of Nontypeable Haemophilus influenzae and Host Cell Transcriptomes Reveals Novel Insights into Host-Pathogen Cross Talk 6(6), e01765– 01715. doi:10.1128/mBio.01765-15.

Bakaletz, L.O., Baker, B.D., Jurcisek, J.A., Harrison, A., Novotny, L.A., Bookwalter, J.E., et al. (2005). Demonstration of type IV pilus expression and a twitching phenotype by Haemophilus influenzae. Infect Immun 73(3), 1635–1643. doi:10.1128/IAI.73.3.1635-1643.2005.

Bell, S.M., Pham, J.N., Rafferty, D.L., and Allerton, J.K. (2016). Antibiotic Susceptibility Testing By the CDS Method (8th Edition) [Online]. Available: http://cdstest.net/wordpress/wp-content/uploads/Antibiotic-Susceptibility-Testing-by-the-CDS-Method-8th-Edition.pdf [Accessed 25/03/2019 2019].

Byun, M.K., Chang, J., Kim, H.J., and Jeong, S.H. (2017). Differences of lung microbiome in patients with clinically stable and exacerbated bronchiectasis. Differences of lung microbiome in patients with clinically stable and exacerbated bronchiectasis 12(8), e0183553. doi:10.1371/journal.pone.0183553.

Coleman, H.N., Daines, D.A., Jarisch, J., and Smith, A.L. (2003). Chemically defined media for growth of Haemophilus influenzae strains. Chemically defined media for growth of Haemophilus influenzae strains 41(9), 4408–4410. doi:10.1128/JCM.41.9.4408-4410.2003.

Connor, T.R., Corander, J., and Hanage, W.P. (2012). Population subdivision and the detection of recombination in non-typable Haemophilus influenzae. Population subdivision and the detection of recombination in non-typable Haemophilus influenzae 158(12), 2958–2964. doi:10.1099/mic.0.063073-0.

De Chiara, M., Hood, D., Muzzi, A., Pickard, D.J., Perkins, T., Pizza, M., et al. (2014). Genome sequencing of disease and carriage isolates of nontypeable Haemophilus influenzae identifies discrete population structure. Genome sequencing of disease and carriage isolates of nontypeable Haemophilus influenzae identifies discrete population structure 111(14), 5439–5444. doi:10.1073/pnas.1403353111.

De Ste Croix, M., Vacca, I., Kwun, M.J., Ralph, J.D., Bentley, S.D., Haigh, R., et al. (2017). Phase-variable methylation and epigenetic regulation by type I restriction-modification systems. Phase-variable methylation and epigenetic regulation by type I restriction-modification systems 41(S1), S3–S15. doi:10.1093/femsre/fux025.

Duell, B.L., Su, Y.C., and Riesbeck, K. (2016). Host-pathogen interactions of nontypeable Haemophilus influenzae: from commensal to pathogen. Host-pathogen interactions of nontypeable Haemophilus influenzae: from commensal to pathogen 590(21), 3840–3853. doi:10.1002/1873-3468.12351.

Erwin, A.L., Nelson, K.L., Mhlanga-Mutangadura, T., Bonthuis, P.J., Geelhood, J.L., Morlin, G., et al. (2005). Characterization of genetic and phenotypic diversity of invasive nontypeable Haemophilus influenzae. Characterization of genetic and phenotypic diversity of invasive nontypeable Haemophilus influenzae 73(9), 5853–5863. doi:10.1128/IAI.73.9.5853-5863.2005.

Erwin, A.L., and Smith, A.L. (2007). Nontypeable Haemophilus influenzae: understanding virulence and commensal behavior. Nontypeable Haemophilus influenzae: understanding virulence and commensal behavior 15(8), 355–362. doi:10.1016/j.tim.2007.06.004.

Fothergill, J.L., Neill, D.R., Loman, N., Winstanley, C., and Kadioglu, A. (2014). Pseudomonas aeruginosa adaptation in the nasopharyngeal reservoir leads to migration and persistence in the lungs. Pseudomonas aeruginosa adaptation in the nasopharyngeal reservoir leads to migration and persistence in the lungs 5, 4780. doi:10.1038/ncomms5780.

Fox, K.L., Dowideit, S.J., Erwin, A.L., Srikhanta, Y.N., Smith, A.L., and Jennings, M.P. (2007). Haemophilus influenzae phasevarions have evolved from type III DNA restriction systems into epigenetic regulators of gene expression. Nucleic Acids Res 35(15), 5242–5252. doi:10.1093/nar/gkm571.

Fung, C., Naughton, S., Turnbull, L., Tingpej, P., Rose, B., Arthur, J., et al. (2010). Gene expression of Pseudomonas aeruginosa in a mucin-containing synthetic growth medium mimicking cystic fibrosis lung sputum. Gene expression of Pseudomonas aeruginosa in a mucin-containing synthetic growth medium mimicking cystic fibrosis lung sputum 59(Pt 9), 1089–1100. doi:10.1099/jmm.0.019984-0.

Galperin, M.Y., Makarova, K.S., Wolf, Y.I., and Koonin, E.V. (2015). Expanded microbial genome coverage and improved protein family annotation in the COG database. Expanded microbial genome coverage and improved protein family annotation in the COG database 43, D261– 269. doi:10.1093/nar/gku1223.

Gibson, L.F., and Khoury, J.T. (1986). Storage and Survival of Bacteria by Ultra-Freeze. Storage and Survival of Bacteria by Ultra-Freeze 3(6), 127–129. doi:10.1111/j.1472-765X.1986.tb01565.x.

Gilsdorf, J.R., Marrs, C.F., and Foxman, B. (2004). Haemophilus influenzae: Genetic Variability and Natural Selection To Identify Virulence Factors. Haemophilus influenzae: Genetic Variability and Natural Selection To Identify Virulence Factors 72(5), 2457–2461. doi:10.1128/iai.72.5.2457-2461.2004.

Grønseth, R., Drengenes, C., Wiker, H.G., Tangedal, S., Xue, Y., Husebø, G.R., et al. (2017). Protected sampling is preferable in bronchoscopic studies of the airway microbiome. Protected sampling is preferable in bronchoscopic studies of the airway microbiome 3(3), 00019–02017. doi:10.1183/23120541.00019-2017.

Hare, K.M., Grimwood, K., Leach, A.J., Smith-Vaughan, H., Torzillo, P.J., Morris, P.S., et al. (2010). Respiratory bacterial pathogens in the nasopharynx and lower airways of Australian indigenous children with bronchiectasis. Respiratory bacterial pathogens in the nasopharynx and lower airways of Australian indigenous children with bronchiectasis 157(6), 1001–1005. doi:10.1016/j.jpeds.2010.06.002.

Holt, L.B. (1962). The growth-factor requirements of Haemopiius influenzae. The growth-factor requirements of Haemopiius influenzae 27, 317–322. doi:10.1099/00221287-27-2-317.

Hood, D.W., Cox, A.D., Gilbert, M., Makepeace, K., Walsh, S., Deadman, M.E., et al. (2001). Identification of a lipopolysaccharide alpha-2,3-sialyltransferase from Haemophilus influenzae. Identification of a lipopolysaccharide alpha-2,3-sialyltransferase from Haemophilus influenzae 39(2), 341–350. doi:doi.org/10.1046/j.1365-2958.2001.02204.x.

Hosking, S.L., Craig, J.E., and High, N.J. (1999). Phase variation of lic1A, lic2A and lic3A in colonization of the nasopharynx, bloodstream and cerebrospinal fluid by Haemophilus influenzae type b. Phase variation of lic1A, lic2A and lic3A in colonization of the nasopharynx, bloodstream and cerebrospinal fluid by Haemophilus influenzae type b 145(11), 3005–3011. doi:10.1099/00221287-145-11-3005.

Humair, P.F., Douet, V., Moran Cadenas, F., Schouls, L.M., Van De Pol, I., and Gern, L. (2007). Molecular identification of bloodmeal source in Ixodes ricinus ticks using 12S rDNA as a genetic marker. Molecular identification of bloodmeal source in Ixodes ricinus ticks using 12S rDNA as a genetic marker 44(5), 869–880. doi:10.1093/jmedent/44.5.869.

Jolley, K.A., and Maiden, M.C. (2010). BIGSdb: Scalable analysis of bacterial genome variation at the population level. BIGSdb: Scalable analysis of bacterial genome variation at the population level 11, 595. doi:10.1186/1471-2105-11-595.

Jurcisek, J.A., Bookwalter, J.E., Baker, B.D., Fernandez, S., Novotny, L.A., Munson, R.S., Jr., et al. (2007). The PilA protein of non-typeable Haemophilus influenzae plays a role in biofilm formation, adherence to epithelial cells and colonization of the mammalian upper respiratory tract. The PilA protein of non-typeable Haemophilus influenzae plays a role in biofilm formation, adherence to epithelial cells and colonization of the mammalian upper respiratory tract 65(5), 1288–1299. doi:10.1111/j.1365-2958.2007.05864.x.

Kasi, A.S., Neubauer, C., Kato, R.M., and Newman, D.K. (2017). Bacterial Growth Rate In Cystic Fibrosis Pulmonary Exacerbation. Bacterial Growth Rate In Cystic Fibrosis Pulmonary Exacerbation 195. doi:10.1164/ajrccm-conference.2017.195.1_MeetingAbstracts.A4856.

Lim, Y.W., Schmieder, R., Haynes, M., Willner, D., Furlan, M., Youle, M., et al. (2013). Metagenomics and metatranscriptomics: windows on CF-associated viral and microbial communities. Metagenomics and metatranscriptomics: windows on CF-associated viral and microbial communities 12(2), 154–164. doi:10.1016/j.jcf.2012.07.009.

Mandrell, R.E., McLaughlin, R., Aba Kwaik, Y., Lesse, A., Yamasaki, R., Gibson, B., et al. (1992). Lipooligosaccharides (LOS) of some Haemophilus species mimic human glycosphingolipids, and some LOS are sialylated. Lipooligosaccharides (LOS) of some Haemophilus species mimic human glycosphingolipids, and some LOS are sialylated 60(4), 1322–1328.

Martinez-Garcia, M.A., Soler-Cataluna, J.J., Donat Sanz, Y., Catalan Serra, P., Agramunt Lerma, M., Ballestin Vicente, J., et al. (2011). Factors associated with bronchiectasis in patients with COPD. Factors associated with bronchiectasis in patients with COPD 140(5), 1130–1137. doi:10.1378/chest.10-1758.

Moleres, J., Fernandez-Calvet, A., Ehrlich, R.L., Marti, S., Perez-Regidor, L., Euba, B., et al. (2018). Antagonistic Pleiotropy in the Bifunctional Surface Protein FadL (OmpP1) during Adaptation of Haemophilus influenzae to Chronic Lung Infection Associated with Chronic Obstructive Pulmonary Disease. Antagonistic Pleiotropy in the Bifunctional Surface Protein FadL (OmpP1) during Adaptation of Haemophilus influenzae to Chronic Lung Infection Associated with Chronic Obstructive Pulmonary Disease 9(5). doi:10.1128/mBio.01176-18.

Murphy, T.F. (2003). Respiratory infections caused by non-typeable Haemophilus influenzae. Respiratory infections caused by non-typeable Haemophilus influenzae 16(2), 129–134. doi:10.1097/01.aco.0000065079.06965.e0.

Murphy, T.F., and Kirkham, C. (2002). Biofilm formation by nontypeable Haemophilus influenzae: strain variability, outer membrane antigen expression and role of pili. BMC Microbiol 2:7. doi:10.1186/1471-2180-2-7.

Murray, N.E. (2000). Type I restriction systems: sophisticated molecular machines (a legacy of Bertani and Weigle). Type I restriction systems: sophisticated molecular machines (a legacy of Bertani and Weigle) 64(2), 412–434. doi:10.1128/MMBR.64.2.412-434.2000.

Pettigrew, M.M., Ahearn, C.P., Gent, J.F., Kong, Y., Gallo, M.C., Munro, J.B., et al. (2018). Haemophilus influenzae genome evolution during persistence in the human airways in chronic obstructive pulmonary disease. Haemophilus influenzae genome evolution during persistence in the human airways in chronic obstructive pulmonary disease. doi:10.1073/pnas.1719654115.

Poole, J., Foster, E., Chaloner, K., Hunt, J., Jennings, M.P., Bair, T., et al. (2013). Analysis of nontypeable Haemophilus influenzae phase-variable genes during experimental human nasopharyngeal colonization. Analysis of nontypeable Haemophilus influenzae phasevariable genes during experimental human nasopharyngeal colonization 208(5), 720–727. doi:10.1093/infdis/jit240.

Power, P.M., Bentley, S.D., Parkhill, J., Moxon, E.R., and Hood, D.W. (2012). Investigations into genome diversity of Haemophilus influenzae using whole genome sequencing of clinical isolates and laboratory transformants. Bmc Microbiol 12:273. doi:10.1186/1471-2180-12-273.

Price, E.P., Harris, T.M., Spargo, J., Nosworthy, E., Beissbarth, J., Chang, A.B., et al. (2017). Simultaneous identification of Haemophilus influenzae and Haemophilus haemolyticus using real-time PCR. Simultaneous identification of Haemophilus influenzae and Haemophilus haemolyticus using real-time PCR 12, 585–593. doi:10.2217/fmb-2016-0215.

Price, E.P., Sarovich, D.S., Mayo, M., Tuanyok, A., Drees, K.P., Kaestli, M., et al. (2013). Within-host evolution of Burkholderia pseudomallei over a twelve-year chronic carriage infection. Within-host evolution of Burkholderia pseudomallei over a twelve-year chronic carriage infection 4(4):e00388–13(4). doi:10.1128/mBio.00388-13.

Price, E.P., Sarovich, D.S., Nosworthy, E., Beissbarth, J., Marsh, R.L., Pickering, J., et al. (2015). Haemophilus influenzae: using comparative genomics to accurately identify a highly recombinogenic human pathogen. BMC Genomics 16, 641. doi:10.1186/s12864-015-1857-x.

Purcell, P., Jary, H., Perry, A., Perry, J.D., Stewart, C.J., Nelson, A., et al. (2014). Polymicrobial airway bacterial communities in adult bronchiectasis patients. Polymicrobial airway bacterial communities in adult bronchiectasis patients 14, 130. doi:10.1186/1471-2180-14-130.

R Core Team (2014). “R: A language and environment for statistical computing”. (Vienna, Austria: R Foundation for Statistical Computing).

Robinson, M.D., McCarthy, D.J., and Smyth, G.K. (2010). edgeR: a Bioconductor package for differential expression analysis of digital gene expression data. edgeR: a Bioconductor package for differential expression analysis of digital gene expression data 26(1), 139–140. doi:10.1093/bioinformatics/btp616.

Santana, E.A., Harrison, A., Zhang, X.J., Baker, B.D., Kelly, B.J., White, P., et al. (2014). HrrF Is the Fur-Regulated Small RNA in Nontypeable Haemophilus influenzae. HrrF Is the Fur-Regulated Small RNA in Nontypeable Haemophilus influenzae 9(8). doi:10.1371/journal.pone.0105644.

Sarovich, D.S., and Price, E.P. (2014). SPANDx: a genomics pipeline for comparative analysis of large haploid whole genome re-sequencing datasets. SPANDx: a genomics pipeline for comparative analysis of large haploid whole genome re-sequencing datasets 7, 618. doi:10.1186/1756-0500-7-618.

Sikkema, D.J., and Murphy, T.F. (1992). Molecular analysis of the P2 porin protein of nontypeable Haemophilus influenzae. Molecular analysis of the P2 porin protein of nontypeable Haemophilus influenzae 60(12), 5204–5211.

Slack, M.P.E. (2015). A review of the role of Haemophilus influenzae in community-acquired pneumonia. A review of the role of Haemophilus influenzae in community-acquired pneumonia 6, 26–43. doi:10.15172/pneu.2015.6/520.

Slack, M.P.E. (2017). The evidence for non-typeable Haemophilus influenzae as a causative agent of childhood pneumonia. The evidence for non-typeable Haemophilus influenzae as a causative agent of childhood pneumonia 9. doi:10.1186/s41479-017-0033-2.

Smith, E.E., Buckley, D.G., Wu, Z., Saenphimmachak, C., Hoffman, L.R., D’Argenio, D.A., et al. (2006). Genetic adaptation by Pseudomonas aeruginosa to the airways of cystic fibrosis patients. Genetic adaptation by Pseudomonas aeruginosa to the airways of cystic fibrosis patients 103(22), 8487–8492. doi:10.1073/pnas.0602138103.

Sriram, K.B., Cox, A.J., Clancy, R.L., Slack, M.P.E., and Cripps, A.W. (2018). Nontypeable Haemophilus influenzae and chronic obstructive pulmonary disease: a review for clinicians. Nontypeable Haemophilus influenzae and chronic obstructive pulmonary disease: a review for clinicians 44(2), 125–142. doi:10.1080/1040841X.2017.1329274.

Starner, T.D., Zhang, N., Kim, G., Apicella, M.A., and McCray, P.B., Jr. (2006). Haemophilus influenzae forms biofilms on airway epithelia: implications in cystic fibrosis. Haemophilus influenzae forms biofilms on airway epithelia: implications in cystic fibrosis 174(2), 213–220. doi:10.1164/rccm.200509-1459OC.

Swords, W.E. (2012). Nontypeable Haemophilus influenzae biofilms: role in chronic airway infections. Nontypeable Haemophilus influenzae biofilms: role in chronic airway infections 2, 97. doi:10.3389/fcimb.2012.00097.

Tufvesson, E., Bjermer, L., and Ekberg, M. (2015). Patients with chronic obstructive pulmonary disease and chronically colonized with Haemophilus influenzae during stable disease phase have increased airway inflammation. Patients with chronic obstructive pulmonary disease and chronically colonized with Haemophilus influenzae during stable disease phase have increased airway inflammation 10, 881–889. doi:10.2147/Copd.S78748.

Van Eldere, J., Slack, M.P.E., Ladhani, S., and Cripps, A.W. (2014). Non-typeable Haemophilus influenzae, an under-recognised pathogen. Non-typeable Haemophilus influenzae, an under-recognised pathogen 14(12), 1281–1292. doi:10.1016/s1473-3099(14)70734-0.

Vasu, K., and Nagaraja, V. (2013). Diverse functions of restriction-modification systems in addition to cellular defense. Diverse functions of restriction-modification systems in addition to cellular defense 77(1), 53–72. doi:10.1128/MMBR.00044-12.

Viberg, L.T., Sarovich, D.S., Kidd, T.J., Geake, J.B., Bell, S.C., Currie, B.J., et al. (2017). Within-Host Evolution of Burkholderia pseudomallei during Chronic Infection of seven Australasian cystic fibrosis patients. Within-Host Evolution of Burkholderia pseudomallei during Chronic Infection of seven Australasian cystic fibrosis patients 8:e00356–17(2). doi:10.1128/mBio.00356-17.

Wong, S.M., and Akerley, B.J. (2012). Genome-scale approaches to identify genes essential for Haemophilus influenzae pathogenesis. Genome-scale approaches to identify genes essential for Haemophilus influenzae pathogenesis 2, 23. doi:10.3389/fcimb.2012.00023.

Wurzel, D.F., Marchant, J.M., Yerkovich, S.T., Upham, J.W., Petsky, H.L., Smith-Vaughan, H., et al. (2016). Protracted Bacterial Bronchitis in Children: Natural History and Risk Factors for Bronchiectasis. Protracted Bacterial Bronchitis in Children: Natural History and Risk Factors for Bronchiectasis 150(5), 1101–1108. doi:10.1016/j.chest.2016.06.030.

Yang, L., Haagensen, J.A., Jelsbak, L., Johansen, H.K., Sternberg, C., Hoiby, N., et al. (2008). In situ growth rates and biofilm development of Pseudomonas aeruginosa populations in chronic lung infections. In situ growth rates and biofilm development of Pseudomonas aeruginosa populations in chronic lung infections 190(8), 2767–2776. doi:10.1128/JB.01581-07.

Zaleski, P., Wojciechowski, M., and Piekarowicz, A. (2005). The role of Dam methylation in phase variation of Haemophilus influenzae genes involved in defence against phage infection. The role of Dam methylation in phase variation of Haemophilus influenzae genes involved in defence against phage infection 151(Pt 10), 3361–3369. doi:10.1099/mic.0.28184-0.

