## Supplementary Text 1 for "Molecular signatures of non-typeable *Haemophilus influenzae* lung adaptation in paediatric chronic lung disease"

### Supplementary Methods

**Whole-Genome Sequencing and Comparative Genomics.** To identify isogenic NP-BAL pairs from each patient, we first performed *in silico* multilocus sequence typing (MLST) on all isolates using the BIGSdb tool (Jolley and Maiden, 2010), which is integrated into the PubMLST database, (https://pubmlst.org/hinfluenzae/) to confirm NP-BAL pairs from each patient had matching sequence types (STs). Isolate data, including novel alleles and STs, were added to this database. To further characterise the genetic diversity and to rule out potential ST homoplasy, the latter of which can confound strain relatedness (De Smet et al., 2015; Aziz et al., 2017), comparative genomic analysis of patient isolates was also carried out using a representative global dataset (Price et al., 2017) of 157 *H. influenzae* strains. SPANDx v3.2 (Sarovich and Price, 2014) (default settings) was used to identify robust single-nucleotide polymorphisms (SNPs) and insertions-deletions (indels) among the isolates based on read mapping to the closed NTHi reference 86-028NP (GenBank: CP000057.2). A maximum parsimony (MP) phylogenetic tree was constructed using PAUP* (v4.0a151) (Swofford, 2001) and visualised using iTOL (Letunic and Bork, 2016). This analysis allowed us to select a diverse set of paired NP-BAL isolates for the current study, and to rule out niche clustering of isolates from NP or lung environments. Based on this phylogeny, we selected 12 paired NP-BAL isolates from 11 patients for further investigation; two genetically distinct NP-BAL pairs were obtained from one patient (designated 60373_P1 and 60373_P2).

Draft genomes were assembled using MGAP v1.0 (Sarovich, 2017) and annotated using Prokka v1.13 (Seemann, 2014) with 86-028NP as the reference. Genetic variant annotation and functional effect prediction were performed using SnpEff v4.1 (Cingolani et al., 2012). The

BEDtools (Quinlan, 2014) output from SPANDx was used to detect large-scale deletions or copy number variants between isogenic strains. To identify chromosome rearrangements, draft genome contigs were first reordered relative to 86-028NP, followed by alignment of the BAL and NP genomes using Mauve v2.4.0 (Darling et al., 2004). Visualisation of alignments and manual inspection of genetic variants were performed using IGV v2.4.14 (Thorvaldsdottir et al., 2013) and Tablet v1.17.08.17 (Milne et al., 2013). Multiple sequence alignments were performed using UGENE v1.31 (Okonechnikov et al., 2012).

**NTHi Liquid Media Growth Conditions for RNA Harvest.** To create LCM, 23 g of special peptone, 1 g starch and 5 g NaCl (Oxoid) was dissolved in 950 mL of ultrapure water and filter-sterilised (0.22 µm Bottle Top Filter; Sigma-Aldrich, Castle Hill, NSW, Australia). Finally, 50 mL of defibrinated horse blood (Oxoid) was added. Lysis of erythrocytes was achieved by gradually heating the media in an 80$^{o}$C water bath for 20 min to release X and V factors. LCM was used for all subsequent liquid culture experiments.

**Viability Counts and Growth Curve Analysis.** Viability counts were performed for all isolates to enable the standardisation of starting inoculum for growth curve analysis. Isolates were cultured on CBA at 24h at 37$^{o}$C with 5% $CO_2$ unless otherwise specified. Lawn growth was harvested and resuspended in sterile 0.9% phosphate-buffered saline (PBS) in Nunc round-bottom culture tubes (Thermo Fisher Scientific, Scoresby, VIC, Australia) to achieve an $OD_{590}$ of 1, as measured by a WPA CO 8000 cell density meter (Biochrom Ltd, Blackburn, VIC, Australia). Duplicate suspensions were created for each isolate to assess viability count reproducibility.

Growth curves for four randomly selected isolates (60295 BAL Hi1, 65290 BAL Hi4, 60068 NP Hi3, and 60373 NP Hi3) were performed to determine the optimal late-log harvest time for

RNA extraction. For each isolate, 4.9 mL LCM aliquots were placed into sterile 50 mL Falcon tubes (Corning Inc., Corning, New York, USA) and inoculated with 100 µL of 5x$10^4$ CFUs. Isolates were incubated in an orbital shaker (BL4620; Bioline Global, Smeaton Grange, NSW, Australia) at 37$^o$C and 200 rpm under aerobic conditions. An hourly sampling of cultures between 6 and 12 hours post-inoculation was performed.

**RNA Extraction.** Isolates were cultured in LCM (final volume 2 mL) using Nunc round-bottom tubes in a closed-cap position to ensure biosafety compliance. LCM was inoculated with ~5x$10^4$ CFU and incubated for 7.5h at 37$^o$C at 200 rpm to achieve late-log growth. All isolates were cultured in duplicate to account for any technical replicability issues. RNA was harvested using TRIzol® on duplicate cultures as per manufacturer's instructions (Ambion, Waltham, USA). Extracts were stored overnight at -80$^o$C to aid in homogenisation. To remove residual DNA, extracts were treated with TURBO™ DNase (Ambion), following the *Rigorous DNase Treatment* instructions.

**RNA Quality Control.** Real-time PCR detection of *siaT* (Aziz et al., 2017) and vertebrate 12S mitochondrial DNA (Humair et al., 2007) was used to verify the removal of NTHi DNA and defibrinated horse blood DNA, respectively. RNA samples were diluted 1:10 in DEPC-treated $H_2O$ (Fisher Biotech, Wembley, Australia) before PCR amplification. Minor modifications were made to each PCR assay as follows. Each *siaT* reaction contained 1× Platinum qPCR SuperMix-UDG (Invitrogen, Scoresby, VIC, Australia), 1 µL of diluted RNA and DEPC-treated $H_2O$ to 10 µL. All PCRs were carried out in duplicate on the Rotor-Gene Q instrument (Qiagen, Chadstone Centre, VIC, Australia), with the 'Auto-Gain Optimisation' option enabled (Rotor-Gene Q software v2.3.1.49). Thermocycling conditions were 95$^o$C for 120 sec, followed by 45 cycles of 95$^o$C for 10 sec and 60$^o$C for 10 sec. *H. influenzae* ATCC® 49247™ DNA was used as positive control. Each vertebrate 12S reaction contained primers at a final concentration of 0.2 µM, 1x Platinum SYBR Green qPCR SuperMix-UDG (Invitrogen), 1 µL of RNA sample and DEPC-treated $H_2O$ to 10 µL. Thermocycling was carried out as follows: 95°C for 120 sec, followed by 45 cycles of 95°C for 10 sec, 52°C for 10 sec, 72°C for 10 sec. Human DNA (~1 ng/µL) and 1:1000 diluted LCM (~0.08 ng/µL horse blood DNA) were included as positive controls. No-template controls were included in all runs. After confirming DNA removal, the Bioanalyzer 2100 (Agilent Technologies, Mulgrave, VIC, Australia) was used to assess total RNA quality (RIN range: 5 to 9) and purity using the RNA 6000 Nano Kit (Agilent) as per manufacturer's instructions.

**Differential Expression (DE) Analysis.** RNA-seq was carried out at the Australian Genome Research Facility (Melbourne, Australia) using the Illumina HiSeq 2500 platform, with all libraries pooled in a single lane. 16S and 23S ribosomal RNA was depleted by treatment with the Ribo-Zero rRNA Removal Kit (Bacteria) (Illumina) followed by 100 bp paired-end, stranded library construction using the TruSeq Stranded messenger RNA Kit (Illumina). In total, 24 isolates were sequenced in duplicate, generating 48 transcriptomes. Read quality was assessed using FastQC (v0.10.1) (Andrews, 2010) and summarised with MultiQC (v1.0) (Ewels et al., 2016). Read trimming and filtering was performed using Trimmomatic (v0.33) (Bolger et al., 2014) with the following parameters: leading = 3, trailing = 3, sliding window = 4:15, minimum length = 36. Alignment of reads to 86-028NP was performed using Bowtie2 (v2.1.0) (Langmead and Salzberg, 2012) with default parameters. Reads were sorted and filtered (mapping quality ≥ 1) with SAMtools (v1.2) (Li et al., 2009). Quantification was performed using HTSeq (v0.9.0) (Anders et al., 2015) with the following parameters: -m intersection-nonempty, -s reverse.

Based on previous studies (Baddal et al., 2015; Price et al., 2018), a $\log_2$ fold change of ≥1.5 and a false discovery rate-adjusted *p* value of ≤0.01 (Benjamini and Hochberg, 1995) was selected to classify DE genes. Counts of technical replicates were summed prior to convergence analysis to account for any technical variability due to culture and RNA extraction. Due to the limited number of biological replicates for each patient, counts of technical replicates were not summed for within-patient analyses. DE analysis was conducted using edgeR (v3.18.1) (Robinson et al., 2010), implemented in R (v3.4.1) (R Core Team, 2014). For all analyses, DE was determined using the quasi-likelihood method (*glmQLFit* function). As part of quality control, multidimensional scaling analysis was performed using normalised counts with all isolates clustering as expected. For data processing, visualisation and cluster analysis, the R packages dplyr (Wickham et al., 2017), Glimma (Su et al., 2017), ggplot2 (Wickham, 2009) and gplots (Warnes et al., 2009) were used.

### References


Anders, S., Pyl, P.T., and Huber, W. (2015). HTSeq--a Python framework to work with high-throughput sequencing data. *HTSeq--a Python framework to work with high-throughput sequencing data* 31(2), 166-169. doi: 10.1093/bioinformatics/btu638.

Andrews, S. (2010). *FastQC: a quality control tool for high throughput sequence data* [Online]. Available: online at: http://www.bioinformatics.babraham.ac.uk/projects/fastqc [Accessed 2018].

Aziz, A., Sarovich, D.S., Harris, T.M., Kaestli, M., McRobb, E., Mayo, M., et al. (2017). Suspected cases of intracontinental *Burkholderia pseudomallei* sequence type homoplasy resolved using whole-genome sequencing. *Suspected cases of intracontinental Burkholderia pseudomallei sequence type homoplasy resolved using whole-genome sequencing* 3(11). doi: 10.1099/mgen.0.000139.

Baddal, B., Muzzi, A., Censini, S., Calogero, R.A., Torricelli, G., Guidotti, S., et al. (2015). Dual RNA-seq of Nontypeable *Haemophilus influenzae* and Host Cell Transcriptomes Reveals Novel Insights into Host-Pathogen Cross Talk. *Dual RNA-seq of Nontypeable Haemophilus influenzae and Host Cell Transcriptomes Reveals Novel Insights into Host-Pathogen Cross Talk* 6(6), e01765-01715. doi: 10.1128/mBio.01765-15.

Benjamini, Y., and Hochberg, Y. (1995). Controlling the False Discovery Rate: A Practical and Powerful Approach to Multiple Testing. *Controlling the False Discovery Rate: A Practical and Powerful Approach to Multiple Testing* 57(1), 289-300.

Bolger, A.M., Lohse, M., and Usadel, B. (2014). Trimmomatic: a flexible trimmer for Illumina sequence data. *Trimmomatic: a flexible trimmer for Illumina sequence data* 30(15), 2114-2120. doi: 10.1093/bioinformatics/btu170.

Cingolani, P., Platts, A., Wang le, L., Coon, M., Nguyen, T., Wang, L., et al. (2012). A program for annotating and predicting the effects of single nucleotide polymorphisms, SnpEff: SNPs in the genome of *Drosophila melanogaster* strain w1118; iso-2; iso-3. *A program for annotating and predicting the effects of single nucleotide polymorphisms, SnpEff: SNPs in the genome of Drosophila melanogaster strain w1118; iso-2; iso-3* 6(2), 80-92. doi: 10.4161/fly.19695.

Darling, A.C., Mau, B., Blattner, F.R., and Perna, N.T. (2004). Mauve: multiple alignment of conserved genomic sequence with rearrangements. *Mauve: multiple alignment of conserved genomic sequence with rearrangements* 14(7), 1394-1403. doi: 10.1101/gr.2289704.

De Smet, B., Sarovich, D.S., Price, E.P., Mayo, M., Theobald, V., Kham, C., et al. (2015). Whole-genome sequencing confirms that *Burkholderia pseudomallei* multilocus sequence types common to both Cambodia and Australia are due to homoplasy. *Whole-genome sequencing confirms that Burkholderia pseudomallei multilocus sequence types common to both Cambodia and Australia are due to homoplasy* 53(1), 323-326. doi: 10.1128/JCM.02574-14.

Ewels, P., Magnusson, M., Lundin, S., and Kaller, M. (2016). MultiQC: summarize analysis results for multiple tools and samples in a single report. *MultiQC: summarize analysis results for multiple tools and samples in a single report* 32(19), 3047-3048. doi: 10.1093/bioinformatics/btw354.

Humair, P.F., Douet, V., Moran Cadenas, F., Schouls, L.M., Van De Pol, I., and Gern, L. (2007). Molecular identification of bloodmeal source in *Ixodes ricinus* ticks using 12S rDNA as a genetic marker. *Molecular identification of bloodmeal source in Ixodes ricinus ticks using 12S rDNA as a genetic marker* 44(5), 869-880. doi: 10.1093/jmedent/44.5.869.

Jolley, K.A., and Maiden, M.C. (2010). BIGSdb: Scalable analysis of bacterial genome variation at the population level. *BIGSdb: Scalable analysis of bacterial genome variation at the population level* 11, 595. doi: 10.1186/1471-2105-11-595.

Langmead, B., and Salzberg, S.L. (2012). Fast gapped-read alignment with Bowtie 2. *Fast gapped-read alignment with Bowtie 2* 9(4), 357-359. doi: 10.1038/nmeth.1923.

Letunic, I., and Bork, P. (2016). Interactive tree of life (iTOL) v3: an online tool for the display and annotation of phylogenetic and other trees. *Interactive tree of life (iTOL) v3: an online tool for the display and annotation of phylogenetic and other trees* 44(W1)**,** W242-245. doi: 10.1093/nar/gkw290.

Li, H., Handsaker, B., Wysoker, A., Fennell, T., Ruan, J., Homer, N., et al. (2009). The Sequence Alignment/Map format and SAMtools. *The Sequence Alignment/Map format and SAMtools* 25(16)**,** 2078-2079. doi: 10.1093/bioinformatics/btp352.

Milne, I., Stephen, G., Bayer, M., Cock, P.J.A., Pritchard, L., Cardle, L., et al. (2013). Using Tablet for visual exploration of second-generation sequencing data. *Brief Bioinform* 14(2)**,** 193-202. doi: 10.1093/bib/bbs012.

Okonechnikov, K., Golosova, O., Fursov, M., and team, U. (2012). Unipro UGENE: a unified bioinformatics toolkit. *Unipro UGENE: a unified bioinformatics toolkit* 28(8)**,** 1166-1167. doi: 10.1093/bioinformatics/bts091.

Price, E.P., Harris, T.M., Spargo, J., Nosworthy, E., Beissbarth, J., Chang, A.B., et al. (2017). Simultaneous identification of *Haemophilus influenzae* and *Haemophilus haemolyticus* using real-time PCR. *Simultaneous identification of Haemophilus influenzae and Haemophilus haemolyticus using real-time PCR* 12**,** 585-593. doi: 10.2217/fmb-2016-0215.

Price, E.P., Viberg, L.T., Kidd, T.J., Bell, S.C., Currie, B.J., and Sarovich, D.S. (2018). Transcriptomic analysis of longitudinal *Burkholderia pseudomallei* infecting the cystic fibrosis lung. *Transcriptomic analysis of longitudinal Burkholderia pseudomallei infecting the cystic fibrosis lung*. doi: 10.1099/mgen.0.000194.

Quinlan, A.R. (2014). BEDTools: The Swiss-Army Tool for Genome Feature Analysis. *BEDTools: The Swiss-Army Tool for Genome Feature Analysis* 47**,** 11 12 11-34. doi: 10.1002/0471250953.bi1112s47.

R Core Team (2014). "R: A language and environment for statistical computing". (Vienna, Austria: R Foundation for Statistical Computing).

Robinson, M.D., McCarthy, D.J., and Smyth, G.K. (2010). edgeR: a Bioconductor package for differential expression analysis of digital gene expression data. *edgeR: a Bioconductor package for differential expression analysis of digital gene expression data* 26(1)**,** 139-140. doi: 10.1093/bioinformatics/btp616.

Sarovich, D. (2017). "MGAP - Microbial Genome Assembler Pipeline". Zenodo).

Sarovich, D.S., and Price, E.P. (2014). SPANDx: a genomics pipeline for comparative analysis of large haploid whole genome re-sequencing datasets. *SPANDx: a genomics pipeline for comparative analysis of large haploid whole genome re-sequencing datasets* 7**,** 618. doi: 10.1186/1756-0500-7-618.

Seemann, T. (2014). Prokka: rapid prokaryotic genome annotation. *Prokka: rapid prokaryotic genome annotation* 30(14)**,** 2068-2069. doi: 10.1093/bioinformatics/btu153.

Su, S., Law, C.W., Ah-Cann, C., Asselin-Labat, M.L., Blewitt, M.E., and Ritchie, M.E. (2017). Glimma: interactive graphics for gene expression analysis. *Glimma: interactive graphics for gene expression analysis* 33(13)**,** 2050-2052. doi: 10.1093/bioinformatics/btx094.

Swofford, D.L. (2001). *PAUP*: Phylogenetic Analysis Using Parsimony (and other methods).* Sinauer Associates.

Thorvaldsdottir, H., Robinson, J.T., and Mesirov, J.P. (2013). Integrative Genomics Viewer (IGV): high-performance genomics data visualization and exploration. *Integrative Genomics Viewer (IGV): high-performance genomics data visualization and exploration* 14(2)**,** 178-192. doi: 10.1093/bib/bbs017.

Warnes, G.R., Bolker, B., Bonebakker, L., Gentleman, R., Huber, W., Liaw, A., et al. (2009). "gplots: Various R programming tools for plotting data", in: *R package version.*).

Wickham, H. (2009). *Ggplot2 : elegant graphics for data analysis.* New York: Springer.

Wickham, H., Francois, R., Henry, L., and Müller, K. (2017). *dplyr: A Grammar of Data Manipulation. R package version 0.7.4* [Online]. Available: https://CRAN.R-project.org/package=dplyr [Accessed].
